# CBAM-enhanced MaxViT-Tiny for gastrointestinal endoscopy image classification: Robustness, explainability, and external validation

**DOI:** 10.64898/2026.10.20.760563

**Authors:** Al Sadek Prince, Israt Fatema Shorna, Md Mehedi Hasan, Mohammad Ahanaf Akif, Aziz Misher Mishu, Gazi Mehraj Sakib

## Abstract

Automated classification of the huge number of images generated during gastrointestinal (GI) endoscopy is required for reliable and interpretable diagnosis, but there is concern about the vulnerability of deep models to adversarial perturbations. In the current study, a MaxViT-Tiny backbone with a Convolutional Block Attention Module (CBAM) head is used for 8-class classification of the Kvasir endoscopic images. After deleting duplicate images, 3,996 images remained, which were split into a training set (70%), a validation set (15%) and a test set (15%); the model was compared with five backbones using the same CBAM head under three random seeds. The proposed model obtained the highest test accuracy of 94.33%, a macro F1 score of 0.9431, and a macro area under the receiver operating characteristic curve (AUC) of 0.9975, with a mean accuracy of 93.72 ± 1.06% across all seeds. White-box FGSM, PGD, CW-L2, and MI-FGSM attacks greatly reduced the accuracy: after adversarial fine-tuning, the accuracy under FGSM was 74.17%, while the accuracy under PGD and MI-FGSM was below 4%, showing poor cross-attack generalization. A preliminary attack detector obtained an AUC of 0.857. Grad-CAM++, Score-CAM, and GradientSHAP sometimes identified clinically relevant regions, but also sometimes focused on non-anatomical artefacts. High accuracy on clean images, even with attention augmentation, therefore does not imply high accuracy on adversarial images or reliable explanations, and the model needs further verification on independent datasets and multi-centre validation before clinical use.

## Introduction

Gastrointestinal (GI) diseases include conditions that impact the esophagus, stomach, small intestine, and colon, such as esophagitis, ulcerative colitis, polyps, ulcers, and GI cancers. Endoscopy is used extensively for their diagnosis, and in the small bowel wireless capsule endoscopy (WCE) generates a large number of images that require interpretation by trained clinicians. Artificial Intelligence (AI) is therefore attracting increasing attention, and a recent systematic survey showed that the number of publications on deep learning, with Convolutional Neural Networks (CNNs) as the most popular architecture, is steadily rising [1]. AI is also used in other GI settings, for instance tabular data analysis for the early detection of GI disorders [2], anaesthesia management during surgery [3], and measurement of slow waves in the stomach and duodenum [4].

Transfer Learning and Attention Mechanisms (TLAM) and Transformer-based designs have recently been used in Endoscopic Image Analysis (EIA). An attention-guided variant of ResNet (SE + ResNet-H) achieved 97.84% accuracy on five classes of HyperKvasir WCE images [5], and attention-augmented transfer learning has been explored for the eight classes of Kvasir [6]. Models fine-tuned with metaheuristics and tested on Kvasir include hybrid rice optimization with an attention-based LSTM [7] and an improved ShuffleNet [8]. An accuracy of 98.2% has been reported on the eight Kvasir classes [9], and 99.50% on the 4-class WCECCD dataset [10] with transformer-based architectures such as TransfoNet. In addition to classification, deep models have also been used for lesion detection [11, 12] and lesion segmentation [13].

Several issues still need to be addressed. The review in [1] lists open points concerning limited real-time effectiveness, difficulties in hospital integration, lack of interpretability, and lack of labelled data. A second issue is evaluation practice: all the image-based studies above report results on a single public benchmark, typically Kvasir or a variant of it, and split the data at the image level [5, 8–10]. When leakage-aware group-level splits and cross-dataset validation were introduced, all compared methods suffered, and the gap with the reported 97%–99% suggests that these numbers may be too optimistic for real-world use, with the improvement limited to 2% on Kvasir-v2 [14]. Several studies have also suggested the need for multi-centre or prospective validation [10, 14], because the performance of fine-tuned transfer-learning models cannot be guaranteed over different devices, centres, or populations, and they need to be validated on larger and more diverse public datasets. Deployment is also difficult: GIFCOS-DT ran at around 5 frames per second (fps) on an embedded device, compared with about 28 fps in the standard setting [12], and models built on a clean benchmark dataset [14] were affected by illumination changes, blur, reflections and viewpoint variation. One important question that has not been studied in GI imaging is adversarial vulnerability: small, hard-to-detect perturbations can change the predictions of deep networks in medical imaging, which is very important for clinical safety [15, 16].

These observations motivate an efficient, robust, accurate and interpretable diagnosis model for GI images. Lightweight architectures, explainable AI, and hybrid CNN–Transformer architectures [1] have shown potential, and further improvements come from attention mechanisms [5], multimodal fusion [9], ensemble learning [7], and joint image-quality screening and abnormality detection [12]. Model decisions have been supported by visualisation such as Grad-CAM [5, 9], but multi-centre validation has not been addressed in the GI studies above. In general, none of them take generalization and adversarial vulnerability into account. Patient-level group splits (stricter testing), cross-dataset testing and multi-centre validation are also important [10, 14]. This study addresses these gaps by using attention- and transformer-based representations, a transparent, deduplicated and seed-repeated evaluation, and an investigation of adversarial robustness and explanation reliability.

### This work’s contributions

- **MaxViT-Tiny classification framework:** We explore an endoscopy image classification framework based on MaxViT-Tiny with a Convolutional Block Attention Module (CBAM) head on the eight classes of the Kvasir dataset, and compare it with five other backbone networks using the same CBAM head.
- **Comprehensive performance evaluation:** The model is evaluated with accuracy, macro-precision, macro-recall, macro-F1 score and macro-AUC, and tests are repeated with three random seeds to assess stability.
- **Adversarial robustness analysis:** We study the robustness of the model against FGSM, PGD, CW-L2, and MI-FGSM attacks, and the effect of adversarial fine-tuning on classification robustness.
- **Explainability, detection and computational analysis:** We evaluate a preliminary adversarial-image detector, explain model predictions with Grad-CAM++, Score-CAM and GradientSHAP, and report the computational cost.
- **External validation:** The trained model is tested on the external GastroVision dataset to assess generalization beyond Kvasir.

### Literature review

The 14 studies reviewed here are analysed by method and summarised in Table 1.

**Table 1.**
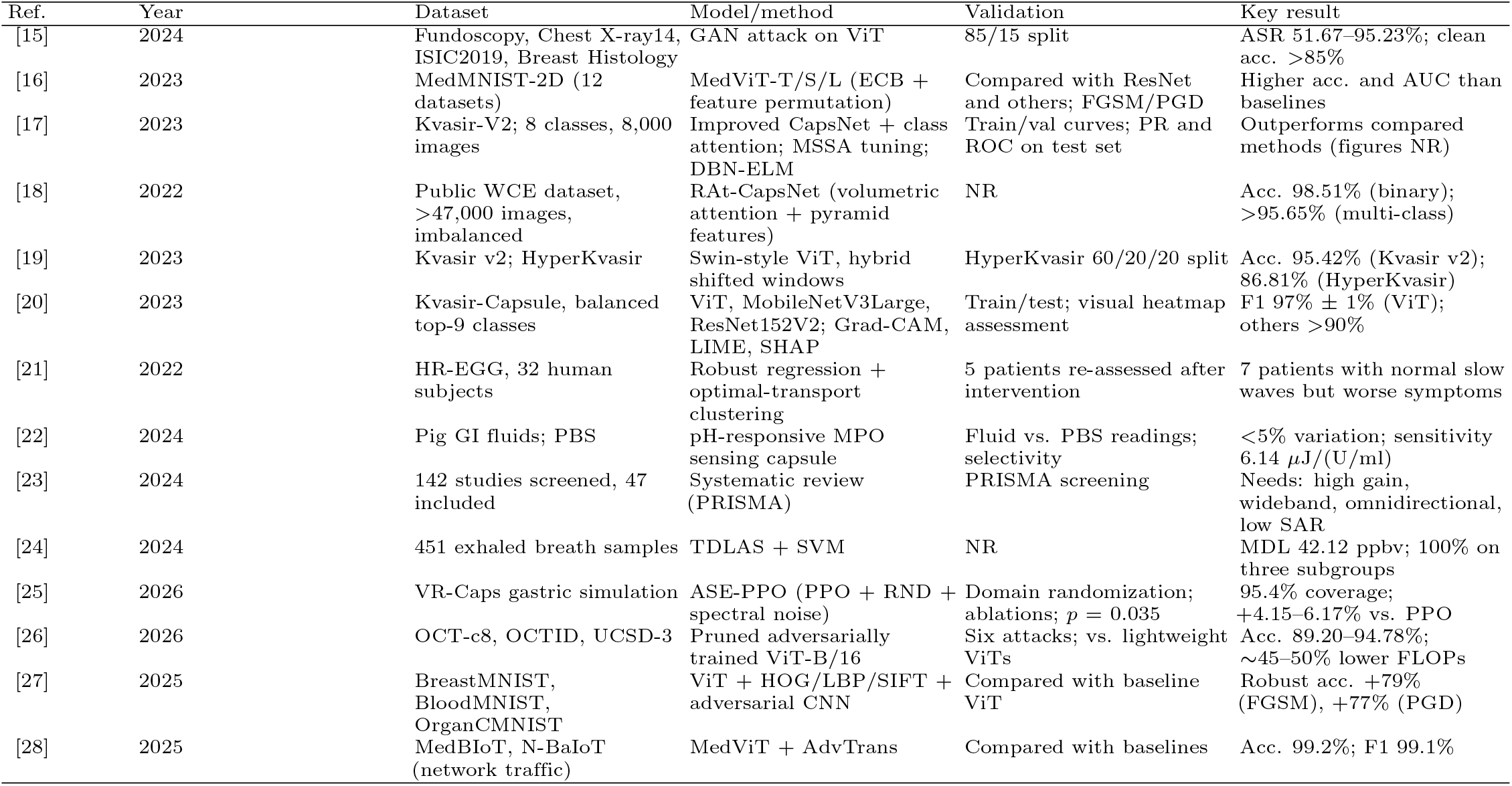
Summary of the studies reviewed in the Literature review section (NR = not reported in the extracted summary; ASR = attack success rate; MDL = minimum detection limit).

| Ref. | Year | Dataset | Model/method | Validation | Key result |
| --- | --- | --- | --- | --- | --- |
| [15] | 2024 | Fundoscopy, Chest X-ray14, ISIC2019, Breast Histology | GAN attack on ViT | 85/15 split | ASR 51.67–95.23%; clean acc. >85% |
| [16] | 2023 | MedMNIST-2D (12 datasets) | MedViT-T/S/L (ECB + feature permutation) | Compared with ResNet and others; FGSM/PGD | Higher acc. and AUC than baselines |
| [17] | 2023 | Kvasir-V2; 8 classes, 8,000 images | Improved CapsNet + class attention; MSSA tuning; DBN-ELM | Train/val curves; PR and ROC on test set | Outperforms compared methods (figures NR) |
| [18] | 2022 | Public WCE dataset, >47,000 images, imbalanced | RAt-CapsNet (volumetric attention + pyramid features) | NR | Acc. 98.51% (binary); >95.65% (multi-class) |
| [19] | 2023 | Kvasir v2; HyperKvasir | Swin-style ViT, hybrid shifted windows | HyperKvasir 60/20/20 split | Acc. 95.42% (Kvasir v2); 86.81% (HyperKvasir) |
| [20] | 2023 | Kvasir-Capsule, balanced top-9 classes | ViT, MobileNetV3Large, ResNet152V2; Grad-CAM, LIME, SHAP | Train/test; visual heatmap assessment | F1 97% ± 1% (ViT); others >90% |
| [21] | 2022 | HR-EGG, 32 human subjects | Robust regression + optimal-transport clustering | 5 patients re-assessed after intervention | 7 patients with normal slow waves but worse symptoms |
| [22] | 2024 | Pig GI fluids; PBS | pH-responsive MPO sensing capsule | Fluid vs. PBS readings; selectivity | <5% variation; sensitivity 6.14 μJ/(U/ml) |
| [23] | 2024 | 142 studies screened, 47 included | Systematic review (PRISMA) | PRISMA screening | Needs: high gain, wideband, omnidirectional, low SAR |
| [24] | 2024 | 451 exhaled breath samples | TDLAS + SVM | NR | MDL 42.12 ppbv; 100% on three subgroups |
| [25] | 2026 | VR-Caps gastric simulation | ASE-PPO (PPO + RND + spectral noise) | Domain randomization; ablations; $p = 0.035$ | 95.4% coverage; +4.15–6.17% vs. PPO |
| [26] | 2026 | OCT-c8, OCTID, UCSD-3 | Pruned adversarially trained ViT-B/16 | Six attacks; vs. lightweight ViTs | Acc. 89.20–94.78%; ~45–50% lower FLOPs |
| [27] | 2025 | BreastMNIST, BloodMNIST, OrganCMNIST | ViT + HOG/LBP/SIFT + adversarial CNN | Compared with baseline ViT | Robust acc. +79% (FGSM), +77% (PGD) |
| [28] | 2025 | MedBioT, N-BaIoT (network traffic) | MedViT + AdvTrans | Compared with baselines | Acc. 99.2%; F1 99.1% |

This section reviews models that classify endoscopic images with Capsule Network and Transformer architectures, followed by physiological sensing and capsule hardware, and adversarial robustness of vision transformers in medical imaging.

Only a few studies classify GI images with non-standard CNNs. On Kvasir-V2 (8,000 images, eight classes), MSSADL-GITDC combined median smoothing, a CapsNet with a class attention layer, salp swarm tuning, and a DBN-ELM classifier, and was reported to outperform the compared methods, but no quantitative details were provided in the source summary [17]. For RAt-CapsNet, the accuracy was 98.51% in the binary case and more than 95.65% in the multi-class case (some pairs of classes have few distinguishing features and low resolution), using more than 47,000 labelled and imbalanced images [18].

Transformer-based classification was explored in [19], where a Swin-style vision transformer with hybrid shifted windows achieved 95.42% accuracy on Kvasir v2 and 86.81% on HyperKvasir (60/20/20 split), indicating that the accuracy depends on the difficulty of the dataset. In [20], the authors used Grad-CAM, Grad-CAM++, LayerCAM, LIME, and SHAP to compare several pretrained networks, including a vision transformer (ViT), MobileNetV3Large, and ResNet152V2, on a balanced subset of the nine largest classes of Kvasir-Capsule. The other two networks had an F1 score of more than 90% ± 1%, while the ViT achieved 97% ± 1%, but the explanation assessment was qualitative, and LIME treated the black borders of the images as important [20].

### Physiological sensing, capsule hardware, and robotic navigation

The second group of studies is based on non-imaging signals or capsule hardware. Robust regression and optimal-transport clustering in a small cohort of 32 human subjects suggested that, although slow-wave characteristics were normal, there was a non-myoelectric aetiology in seven patients with worse symptoms [21]. A single-use ingestible capsule (11 mm × 26 mm) with a chemiluminescent myeloperoxidase (MPO) sensor and a pH-responsive coating was developed to detect inflammation; in pig GI fluids it showed less than 5% variation in readings and a sensitivity of 6.14 *µ*J/(U/ml), but in vivo results have not been reported [22]. A PRISMA review of 47 studies concluded that capsule-to-receiver telemetry links for WCE should have high gain, wide bandwidth, an omnidirectional pattern, and a low specific absorption rate (SAR) [23]. TDLAS combined with SVM classification achieved 100% accuracy for ileus, ascending colon cancer, and cancers with liver metastasis, while no significant correlation was found for GIST, stomach cancer, or rectal cancer, which may be due to the small number of patients [24]. For autonomous capsule navigation, ASE-PPO achieved 95.4% stomach coverage at 120 s of simulation, a 4.15%–6.17% improvement over PPO, but questions about the gap between simulation and the real world, and about the safety of magnetic control, remain open [25].

### Adversarial robustness of vision transformers for medical imaging

A final class of studies concerns adversarial robustness, as transformers are beginning to be used in medical imaging. These works do not use GI data, but are relevant to the issues faced by endoscopic models. A GAN-based attack generator was applied to Fundoscopy, Chest X-ray14, ISIC2019, and Breast Histology data, with attack success rates of 81.36%, 95.23%, 77.75%, and 51.67%, respectively, the lowest being on the smallest dataset [15]. GreenShield, an adversarially trained ViT-B/16 with anti-saturation, was tested on three retinal OCT datasets; it saved 45%–50% of FLOPs, inference time, and carbon footprint while keeping higher robustness to six white-box attacks than other lightweight ViTs [26]. Hy-Deft improves the baseline ViT by up to 79% (FGSM) and 77% (PGD) in robust accuracy on three MedMNIST datasets [27] by combining ViT features with HOG, LBP, and SIFT descriptors and an adversarially trained light CNN. MAViT, which combines MedViT with an adversarial-aware transformer, obtained 99.2% accuracy on MedBIoT and N-BaIoT [28] (an IoT-security application). The hybrid CNN–Transformer MedViT, with an Efficient Convolution Block and feature-statistic permutation, outperformed ResNet on 12 MedMNIST-2D datasets and showed better performance against FGSM and PGD than ResNet [16].

### Summary and research gap

Taken together, the studies show that capsule-network and transformer models achieve high accuracy on endoscopic and capsule endoscopy images [17–20], but comparing their accuracy is difficult because the studies use different evaluation protocols, dataset sizes, image crops, and class distributions. Other diagnostic approaches, such as physiological sensing, breath analysis and capsule hardware, have been tested only in limited numbers of subjects, in animals, with collected fluids, or in simulation [21, 22, 24, 25]. Although powerful yet lightweight vision transformers have been tested on retinal, chest, dermoscopic, histological and MedMNIST images [15, 16, 26–28], no study has examined them on GI endoscopic images. The present work is motivated by this lack of knowledge about how attention-based and transformer-based GI classifiers behave under perturbations, under patient-level validation, and across datasets.

## Materials and methods

### Dataset

The experiments are carried out on the original Kvasir dataset (version 1), which comprises gastrointestinal (GI) endoscopic images (4,000 images, 500 per class) [29]. This is not the 8,000-image Kvasir-v2 release used in [17] and [19]. The dataset consists of eight classes: three anatomical landmarks (normal-z-line, normal-pylorus, normal-cecum), three pathological findings (esophagitis, polyps, ulcerative-colitis), and two classes of endoscopic procedures (dyed-lifted-polyps, dyed-resection-margins). All files were hashed with SHA-256, and duplicate files (bit-by-bit identical) were removed before splitting; the script aborts if two identical files have different labels. After this step, 3,996 images remained (500 per class, except polyps and ulcerative-colitis, which have 498 images each).

### Data partitioning

All images were split into training, validation and test sets with stratified sampling using a fixed split seed of 42, so that class proportions are maintained and all models and training seeds are evaluated on the same 600 test images (Table 2). The validation set was used for checkpoint selection, early stopping, and learning-rate scheduling, and the test set was used only for the final evaluation. The split is at the image level only: patient or procedure identifiers are not included in the dataset, and near-duplicate frames from the same examination cannot be excluded because only byte-identical frames were removed. This is a limitation of the evaluation protocol.

**Table 2.** Number of images per class in each subset.

| Class | Train | Validation | Test | Total |
| --- | --- | --- | --- | --- |
| dyed-lifted-polyps | 350 | 75 | 75 | 500 |
| dyed-resection-margins | 350 | 75 | 75 | 500 |
| esophagitis | 350 | 75 | 75 | 500 |
| normal-cecum | 350 | 75 | 75 | 500 |
| normal-pylorus | 350 | 75 | 75 | 500 |
| normal-z-line | 350 | 75 | 75 | 500 |
| polyps | 348 | 75 | 75 | 498 |
| ulcerative-colitis | 349 | 74 | 75 | 498 |
| Total | 2797 | 599 | 600 | 3996 |

The test set contains 75 images per class.

### Preprocessing and data augmentation

All images were converted to RGB, resized to 256 × 256 pixels, and normalised with the ImageNet channel means (0.485, 0.456, 0.406) and standard deviations (0.229, 0.224, 0.225). On-the-fly augmentation of training images included a random resized crop (scale 0.80–1.0), random horizontal flip, colour jitter (brightness 0.15, contrast 0.15, saturation 0.10) and a random rotation (up to 12^*°*^). Validation and test images were only resized to 256 × 256 and centre-cropped to 224 × 224. Class imbalance was checked automatically: a weighted random sampler is enabled only if the ratio between the largest and smallest training class is ≥ 1.25. The classes are almost balanced (ratio approx. 1.01); therefore, ordinary random shuffling of the training images is used together with class-weighted cross-entropy. Two representative images for each class are displayed in Fig 1.

**Fig 1.**
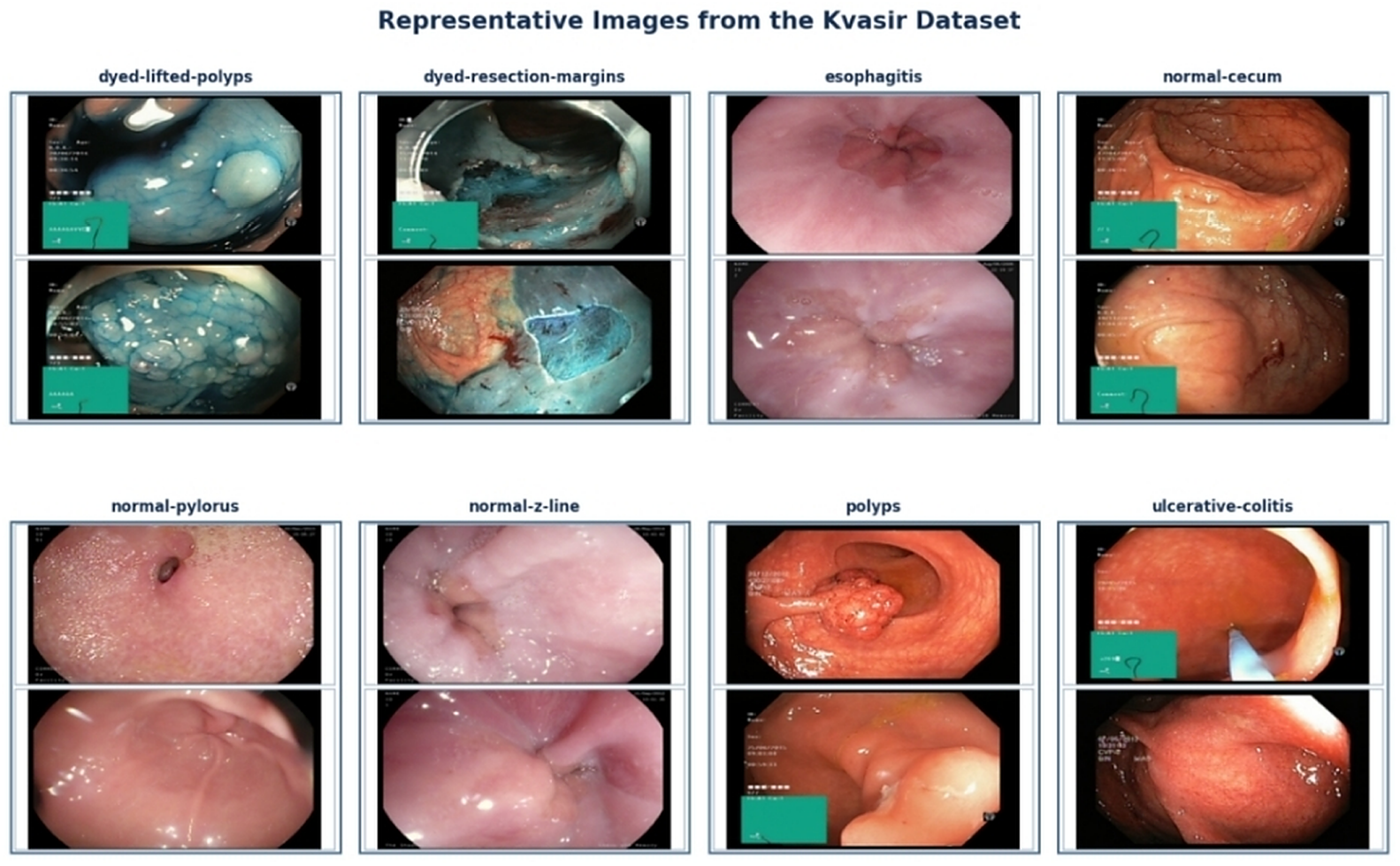
Sample input images from the Kvasir dataset [29].

### Proposed model architecture

The proposed network is composed of two components: the MaxViT-Tiny backbone [30] and a Convolutional Block Attention Module (CBAM) [31] (Fig 2). All the backbones used in this study are compared using the same CBAM head, attached to the final feature map of each backbone. MaxViT-Tiny is a hybrid convolution–transformer network. It is composed of a convolutional stem and four stages of MaxViT blocks, each of which stacks MBConv layers, block (windowed local) self-attention, and grid (dilated global) self-attention, enabling both local and global dependencies to be captured at linear complexity. The backbone was initialised from the “timm” library [32] checkpoint maxvit tiny tf 224.in1k, pretrained on ImageNet-1k, and the original classification head was removed, leaving the last feature map of the backbone (7 *×* 7 *×* 512 for a 224 *×* 224 input) as the output.

**Fig 2.**
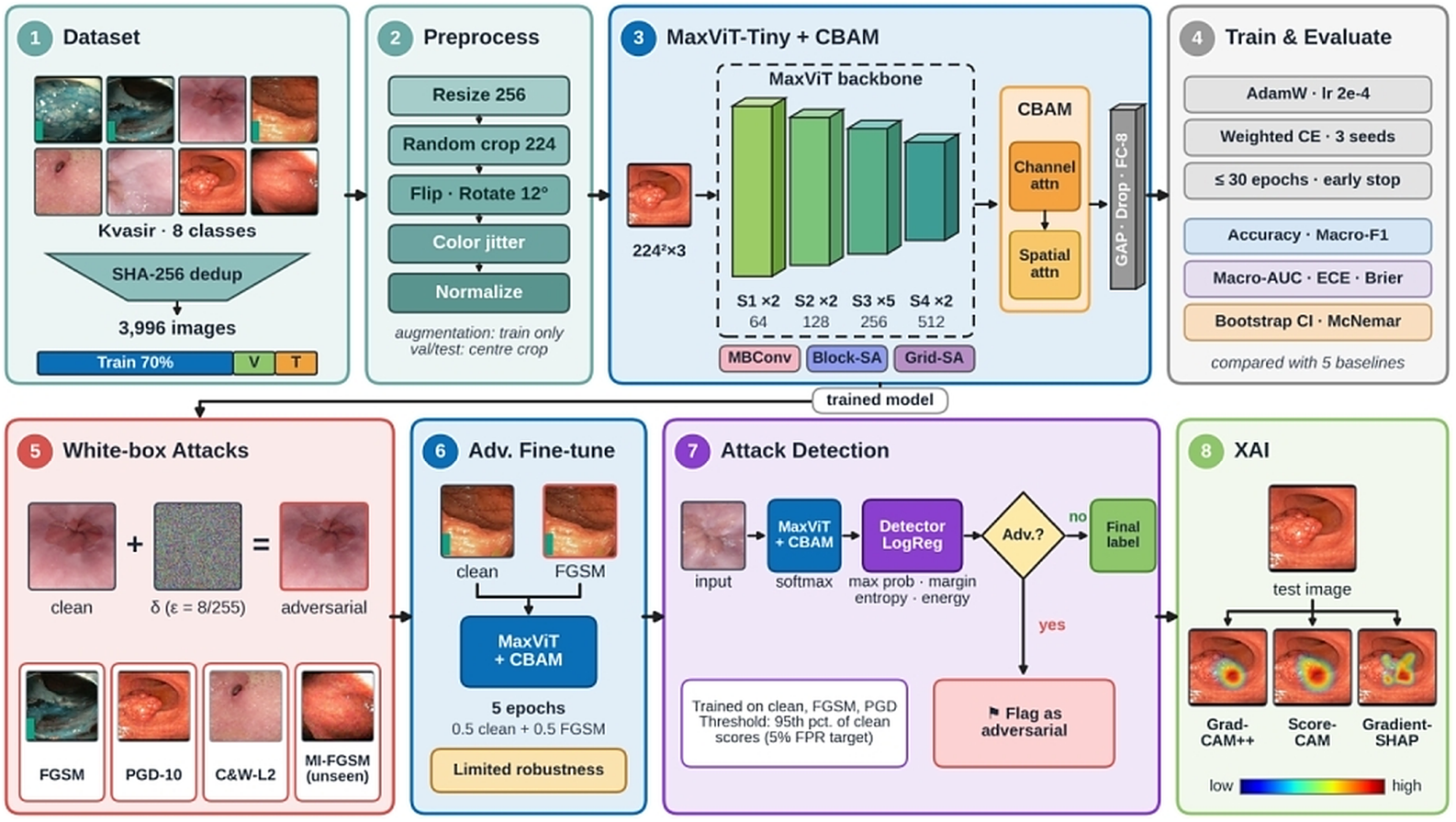
Architecture of the proposed MaxViT-Tiny classifier with a CBAM head.

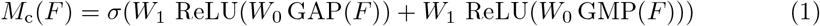

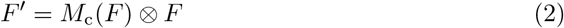

The spatial-attention branch concatenates the channel-wise average and maximum maps of *F*^*′*^, applies a 7 × 7 convolution and a sigmoid gate, and rescales the spatial positions (Eqs 3 and 4):

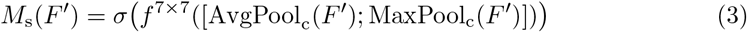

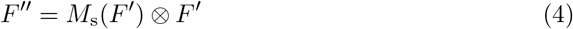

where *σ* is the sigmoid function, *W*_0_ and *W*_1_ are the weights of the shared perceptron, ⊗ is element-wise multiplication with broadcasting, and [· ; ·] is concatenation along the channel axis. The refined feature map *F* ^*′′*^ is reduced by global average pooling, passed through dropout (*p* = 0.2) and a fully connected layer with eight outputs, followed by a softmax.

### Training procedure

The model was trained end-to-end with automatic mixed precision and the AdamW optimiser [33] for a maximum of 30 epochs, with a batch size of 16, an initial learning rate of 2 × 10^−4^, and a weight decay of 10^−4^. The loss function was class-weighted cross-entropy with label smoothing [34] (Eq 5). The main settings are summarised in Table 3.

**Table 3.**
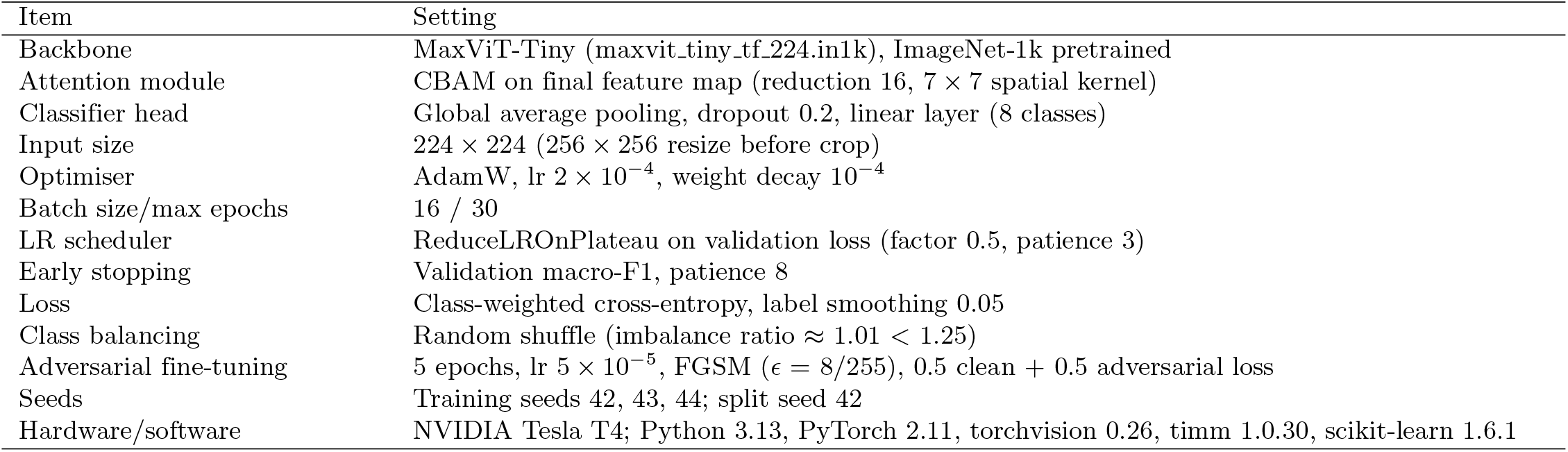
Implementation details and hyperparameters.

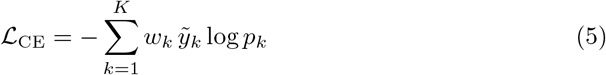

where *K* = 8 is the number of classes, *p*_*k*_ is the predicted probability of class *k*, 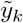 is the smoothed label (*y*_*k*_ is the one-hot label and the smoothing factor is *α* = 0.05), and *w*_*k*_ is the class weight (computed from *n*_*k*_, the number of training images of class *k*, and *n*, the total number of training images). A ReduceLROnPlateau scheduler halved the learning rate if the validation loss did not improve for three epochs. After each training epoch, the checkpoint with the highest validation macro-F1 score was retained, and training was terminated if the validation macro-F1 score did not improve for eight consecutive epochs. In the reported run (seed 44), the best checkpoint was found at epoch 3 (validation macro-F1 0.9533) and training was terminated at epoch 11 (Fig 3). Training was repeated with three random seeds (42, 43 and 44) and the data split was unchanged. For reproducibility, deterministic cuDNN and the same seeds for Python, NumPy and PyTorch were used.

**Fig 3.**
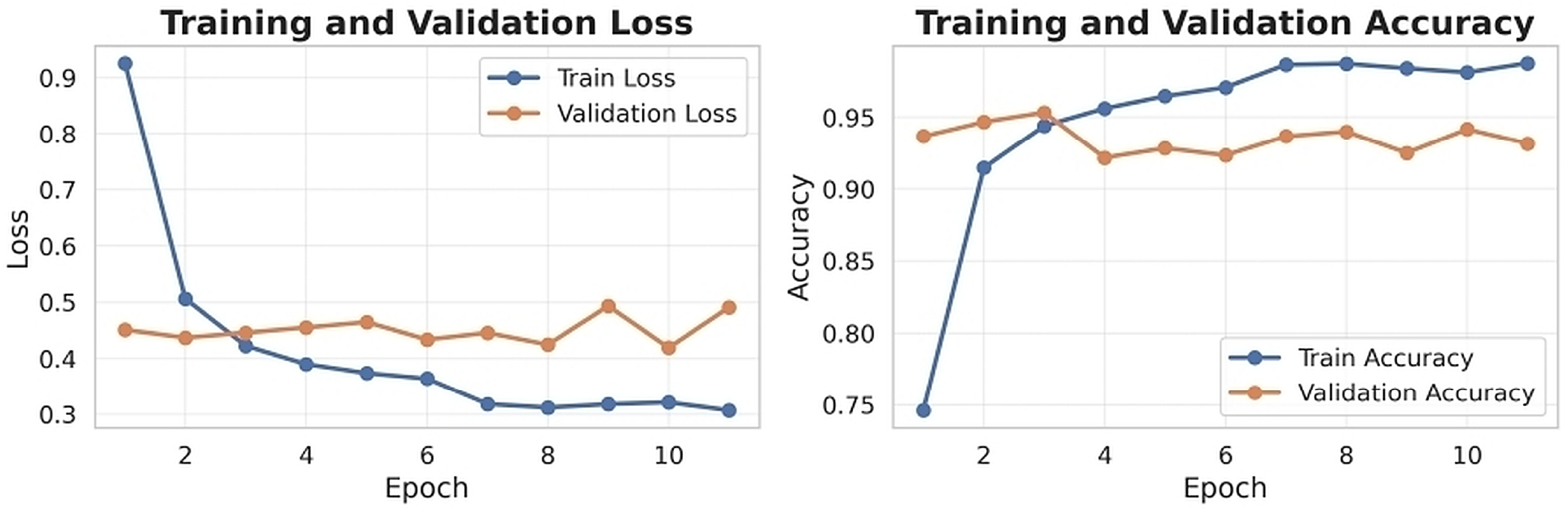
Training and validation loss and accuracy of the proposed model (seed 44).

### Adversarial fine-tuning

To improve robustness, the best clean checkpoint was fine-tuned for five epochs with a mixed clean and adversarial objective (Eq 6), where the adversarial images were generated on the fly with an untargeted FGSM attack [35] (Eq 7):

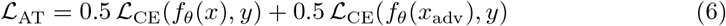

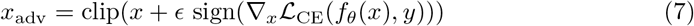

with *ϵ* = 8/255 in pixel space (applied in normalised space) and clip(·) keeping the image in the valid range. The learning rate was reduced to one quarter of the original value (5 × 10^−5^), and the checkpoint with the highest clean validation macro-F1 was retained (Fig 4).

**Fig 4.**
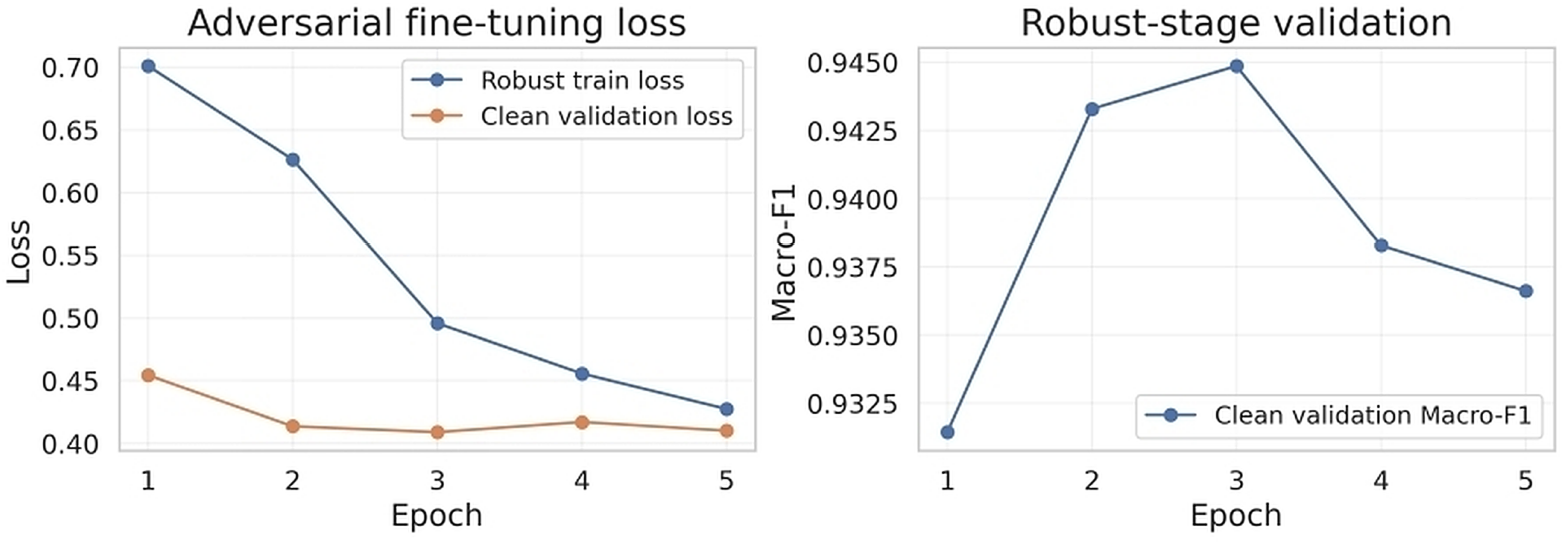
Adversarial fine-tuning: training loss, clean validation loss, and clean validation macro-F1 per epoch.

### Baseline models and comparison protocol

Five models were compared with the proposed model: ConvNeXt-Tiny [36], a hybrid of ConvNeXt-Tiny [36] and Swin-Tiny [37], DenseNet121 [38], GhostNet-100 [39], and MobileNetV3-Large-100 [40]. All models were tested on the same 600 test images, the order of the images in the prediction files stored by the models was verified to be identical, and three different training seeds (42, 43, 44) were used. The main comparison is given for seed 44; consistency across seeds is given as mean ± standard deviation. The same CBAM module as in the proposed model is applied to the final feature map of every baseline, so this is a backbone comparison. The class-balancing strategy differed between models: ConvNeXt-Tiny, DenseNet121, GhostNet, and MobileNetV3 used a weighted random sampler, while MaxViT-Tiny and the ConvNeXt-Tiny + Swin-Tiny hybrid used ordinary random shuffling.

### Evaluation metrics

Performance was measured on the test set with the following metrics. Let TP, TN, FP, and FN denote the one-vs-rest counts for each class, and let *K* = 8 be the number of classes.

Accuracy is the fraction of correctly classified test images. The class-wise metrics are defined as follows:

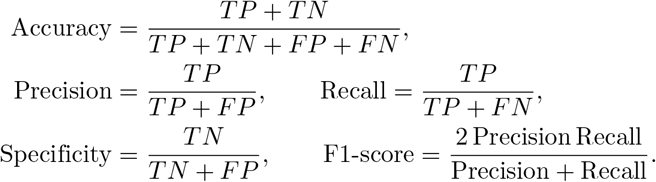

Class-wise values are reported together with macro averages, computed as the unweighted mean over classes, and weighted averages. For any metric *M*, the macro average is:

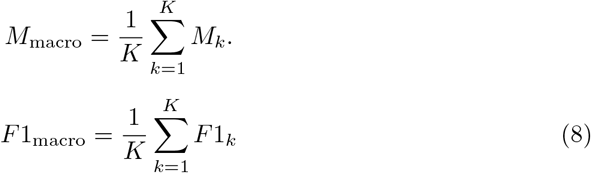

Cohen’s kappa [41] is reported as a supplementary measure of agreement.

The area under the ROC curve (AUC) is computed one-vs-rest for each class and macro-averaged.

The confusion matrix (raw counts and row-normalised) is used to analyse the errors between classes.

Statistical significance: McNemar’s exact test [42] was applied to the paired test predictions of two models, using the discordant pairs, where *b* is the number of images classified correctly only by the first model and *c* is the number classified correctly only by the second (Eq 9). Each comparison was performed separately per seed (five baselines × three seeds = 15 tests), and two-sided exact *p*-values are reported.

Bonferroni-corrected thresholds are given both within a seed (0.05/5 = 0.01) and across all 15 comparisons (0.05/15 ≈ 0.0033), and the Holm step-down procedure is used as a less conservative check. Because the three seeds share the same split and test images, the tests are not independent replications, and McNemar’s test addresses differences between the trained models on this test set only, not the variability due to training randomness or to a different patient sample.

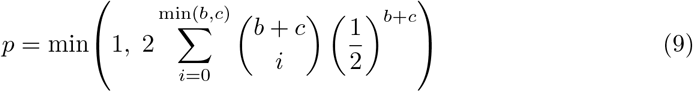

Computational cost: number of parameters, model size, multiply–accumulate operations (MACs) and FLOPs for one 224 × 224 image (thop), and the inference latency and throughput on the GPU for a single image (batch size 1) after 20 warm-up runs, averaged over 100 runs.

### Adversarial robustness and attack detection

Robustness was evaluated on the complete test set with white-box, untargeted attacks against both the clean-trained and the adversarially fine-tuned checkpoint. FGSM followed Eq (7). PGD [43] iterates the signed-gradient step with step size *α* and projects the result onto the *ϵ*-ball *B*_*ϵ*_(*x*) around the input (Eq 10):

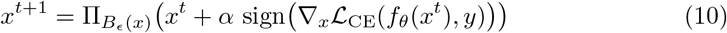

MI-FGSM [44] accumulates the *L*_1_-normalised gradient with decay factor *µ* (Eq 11) and updates the image with the sign of the accumulated momentum (Eq 12):

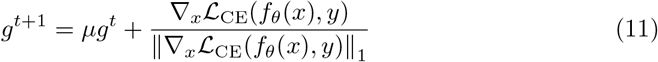

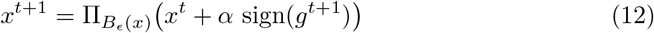

The CW-L2 attack [45] minimises the perturbation norm plus a margin loss on the logits *Z*, with confidence *κ* (Eq 13):

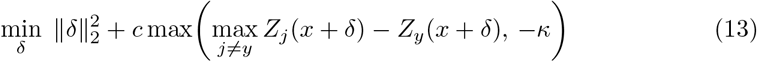

FGSM, PGD, and MI-FGSM used *ϵ* = 8/255 in normalised pixel space; PGD used 10 steps (*α* = 2/255) and a random start, while MI-FGSM used 10 steps (*α* = 2/255) and *µ* = 0.9 and was not used during fine-tuning. CW-L2 used 30 optimisation steps, *c* = 1, and *κ* = 0. We report the accuracy under attack (robust accuracy) and the attack success rate (ASR), defined as the fraction of originally correct predictions that were flipped by the attack (Eq 14):

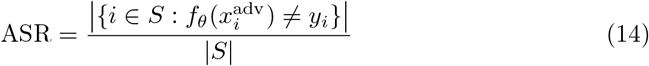

where *S* is the set of test images correctly classified before the attack.

As a preliminary defence, a logistic-regression attack detector was trained on four statistics of the softmax output, computed from clean, FGSM, and PGD images drawn from 50 validation images per class (Eq 15):

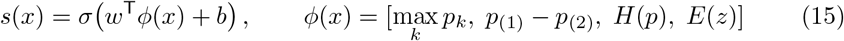

where *p* is the softmax vector, *p*_(1)_ and *p*_(2)_ are its two largest entries, *z* are the logits, and the entropy *H* and energy score *E* [46] are given by Eq (16):

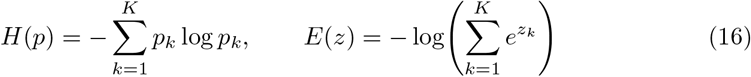

An image was flagged as adversarial when *s*(*x*) exceeded a threshold *τ*, set to the 95th percentile of the scores of 80 clean validation images (10 per class), targeting a 5% false-positive rate. The detector was evaluated on the clean and attacked test images.

### Explainability

The model decisions were examined using three post hoc methods: Grad-CAM++ [47] on the final convolutional layer, Score-CAM [48] on the preceding, higher-resolution convolutional layer, and GradientSHAP [49] (Captum [50], zero and low-noise Gaussian baselines, 20 samples, noise standard deviation 0.09). The first correctly classified test image of each class was visualised, and the explanation was computed for the predicted class. These maps are qualitative in nature.

### External validation dataset

To check the generalization beyond Kvasir, the trained model was also tested on an external dataset, GastroVision [51]. GastroVision is an open-access gastrointestinal endoscopy dataset containing 8,000 images of 27 classes, including anatomical landmarks, pathological findings, polyp-removal cases, and normal cases. The images were collected at Bærum Hospital (Norway) and Karolinska University Hospital (Sweden), and were labelled and reviewed by experienced endoscopists. The data were downloaded from the Kaggle mirror “orvile/gastrovision-gastrointestinal-disease-detection” of the dataset hosted on the Open Science Framework (https://osf.io/84e7f/). No GastroVision image was used for training, validation, hyperparameter selection, or checkpoint selection; the model was trained only on Kvasir.

The label sets of GastroVision and Kvasir are different. Only classes that clinically match a Kvasir label were kept, and matching was performed on the GastroVision folder names (Table 11). Seven of the eight Kvasir classes have a counterpart in GastroVision. GastroVision classes without a Kvasir equivalent (e.g., gastric polyps, resected polyps, and mucosal inflammation of the large bowel) were not included, and ulcerative-colitis, which has no counterpart, was excluded. The mapping was not re-checked by a clinician.

All selected files were hashed with SHA-256 before the evaluation. A single repeated image in the external data was removed (byte-identical), and no external image duplicated an image in Kvasir, resulting in 2,149 external images. The classes are imbalanced (819 vs. 107 images). The Kvasir data were collected at Vestre Viken Health Trust, including Bærum Hospital [29], and no patient identifiers are available for either dataset. Therefore, near-duplicate frames or patient overlap between Kvasir and GastroVision cannot be excluded, and the evaluation is a cross-dataset test and not a fully independent multi-centre validation.

### Preprocessing and protocol for the external evaluation

As for the Kvasir validation and test images (Section 3.3), the external images were converted to RGB, resized to 256 × 256, centre-cropped to 224 × 224, and normalised with the ImageNet means and standard deviations, without augmentation. The seed-44 MaxViT-Tiny checkpoint, trained on Kvasir only, was used in inference mode with no weights adjusted and a batch size of 32. Since the model has eight outputs and the external data have seven classes, the softmax probabilities *p* were renormalised over the set *S* of the seven classes (Eq 17), and all metrics were computed on this seven-class problem, using the arg-max of the renormalised probabilities as the predicted class:

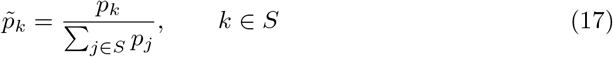

The more stringent eight-way arg-max was also assessed, where a prediction of ulcerative-colitis is counted as an error. As in Section 3.8, accuracy, balanced accuracy, macro precision and recall, macro AUC (one-vs-rest), MCC [52], Cohen’s kappa and ECE [53] (15 bins) were calculated. 2,000 class-stratified bootstrap resamples were used to obtain the 95% confidence interval of the macro F1-score. The results on the Kvasir test set and the external results are compared in one table (Table 12).

## Results and discussion

### Comparison with baseline models

Table 4 shows the comparison of the proposed MaxViT-Tiny with the five baseline networks on the held-out test set (seed 44). MaxViT-Tiny obtained the best results on all metrics: an accuracy of 94.33%, a macro F1-score of 0.9431, and a macro AUC of 0.9975. The nearest baseline was the ConvNeXt-Tiny + Swin-Tiny hybrid (93.33%), 1.00 percentage point lower. The other baselines (ConvNeXt-Tiny, DenseNet121, GhostNet, and MobileNetV3) were 3.50–12.16 percentage points lower than the proposed model. Precision, recall, and F1-score are macro-averaged over the eight classes.

**Table 4.**
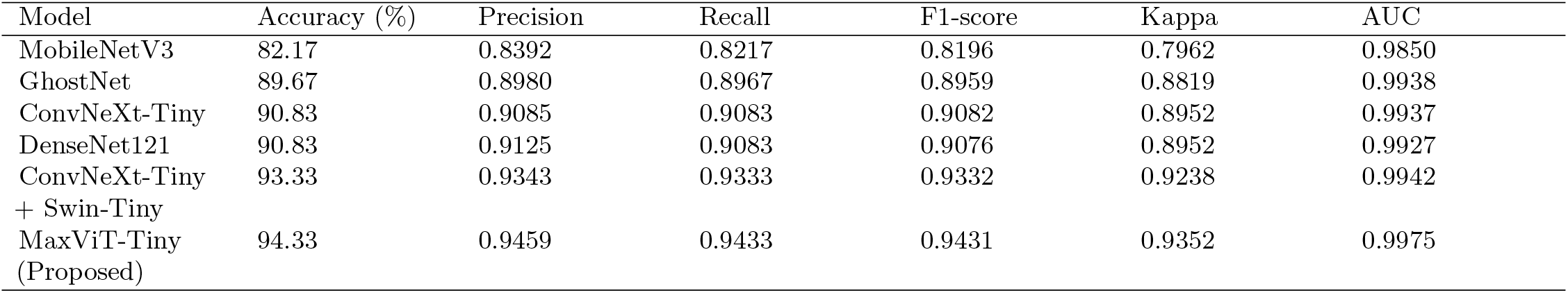
Performance comparison of the proposed MaxViT-Tiny model and baseline models on the test set (seed 44).

### Class-wise performance of the proposed model

The class-wise results of the proposed model are shown in Table 5. The normal-cecum, normal-pylorus, and ulcerative-colitis classes were classified almost perfectly (F1 = 0.9934, 0.9868, and 0.9801, respectively), and every class had an AUC of at least 0.9924. The weakest classes were esophagitis (F1 = 0.8777) and normal-z-line (F1 = 0.8944). These two classes are mainly confused with each other: 14 of the 75 esophagitis images were predicted as normal-z-line, which lowered the recall of esophagitis (0.8133) and the precision of normal-z-line (0.8372). Another source of error was the pair of “dyed” classes: eight dyed-resection-margins images were predicted as dyed-lifted-polyps, and four dyed-lifted-polyps images as dyed-resection-margins.

**Table 5.** Class-wise results of the proposed MaxViT-Tiny (seed 44).

| Class | Precision | Recall | F1-score | Specificity | AUC | Support |
| --- | --- | --- | --- | --- | --- | --- |
| dyed-lifted-polyps | 0.8987 | 0.9467 | 0.9221 | 0.9848 | 0.9974 | 75 |
| dyed-resection-margins | 0.9437 | 0.8933 | 0.9178 | 0.9924 | 0.9975 | 75 |
| esophagitis | 0.9531 | 0.8133 | 0.8777 | 0.9943 | 0.9928 | 75 |
| normal-cecum | 0.9868 | 1.0000 | 0.9934 | 0.9981 | 1.0000 | 75 |
| normal-pylorus | 0.9740 | 1.0000 | 0.9868 | 0.9962 | 1.0000 | 75 |
| normal-z-line | 0.8372 | 0.9600 | 0.8944 | 0.9733 | 0.9924 | 75 |
| polyps | 1.0000 | 0.9467 | 0.9726 | 1.0000 | 0.9998 | 75 |
| ulcerative-colitis | 0.9737 | 0.9867 | 0.9801 | 0.9962 | 0.9998 | 75 |
| Macro average | 0.9459 | 0.9433 | 0.9431 | 0.9919 | 0.9975 | 600 |

Specificity is computed one-vs-rest. Sensitivity equals recall and is therefore not repeated.

### Training behaviour

The training of the proposed model (seed 44) is illustrated in Fig 3. The training loss decreased from 0.93 to 0.31 within 11 epochs, while the training accuracy increased from 74.6% to 98.7%, and the validation loss stayed in the range of 0.42 to 0.49 with the validation accuracy between 92% and 95%. The best validation macro-F1 score (0.9533) was reached at the 3rd epoch, and early stopping ended the training at the 11th epoch. The gap between training and validation accuracy suggests some overfitting, which is to be expected when fine-tuning a pretrained transformer on only about 2,800 images; for this reason, the checkpoint was selected on the validation set and not on the training set. During adversarial fine-tuning, the robust training loss decreased gradually over the five epochs (Fig 4) without any noticeable change in the clean validation loss (note the zoomed vertical axis in Fig 4).

### Error analysis

Fig 5 shows the row-normalised confusion matrix of the proposed MaxViT-Tiny model. The model obtained a recall of at least 0.81 for each of the eight classes and a mean per-class recall of 0.9433, which is good across anatomical landmarks, pathological findings, and endoscopic procedures. Perfect recall (1.00) was obtained for normal-cecum and normal-pylorus, and high recall for ulcerative-colitis (0.99), normal-z-line (0.96), polyps (0.95), and dyed-lifted-polyps (0.95). Misclassifications did not occur at random but tended to be grouped in clinically and visually close pairs of classes. Esophagitis had the lowest recall (0.81), with 19% of the samples classified as normal-z-line, which is likely explained by the similar appearance of the gastric and oesophageal mucosa, where early inflammatory changes may be subtle at the gastro-oesophageal (GE) junction. Likewise, dyed-resection-margins (0.89) and dyed-lifted-polyps (0.95) were confused in both directions, with 11% and 5% of the samples misassigned, respectively; this may be explained by the similar chromoendoscopic dye-staining properties. All other off-diagonal entries were no more than 0.04, indicating little confusion between the remaining classes.

**Fig 5.**
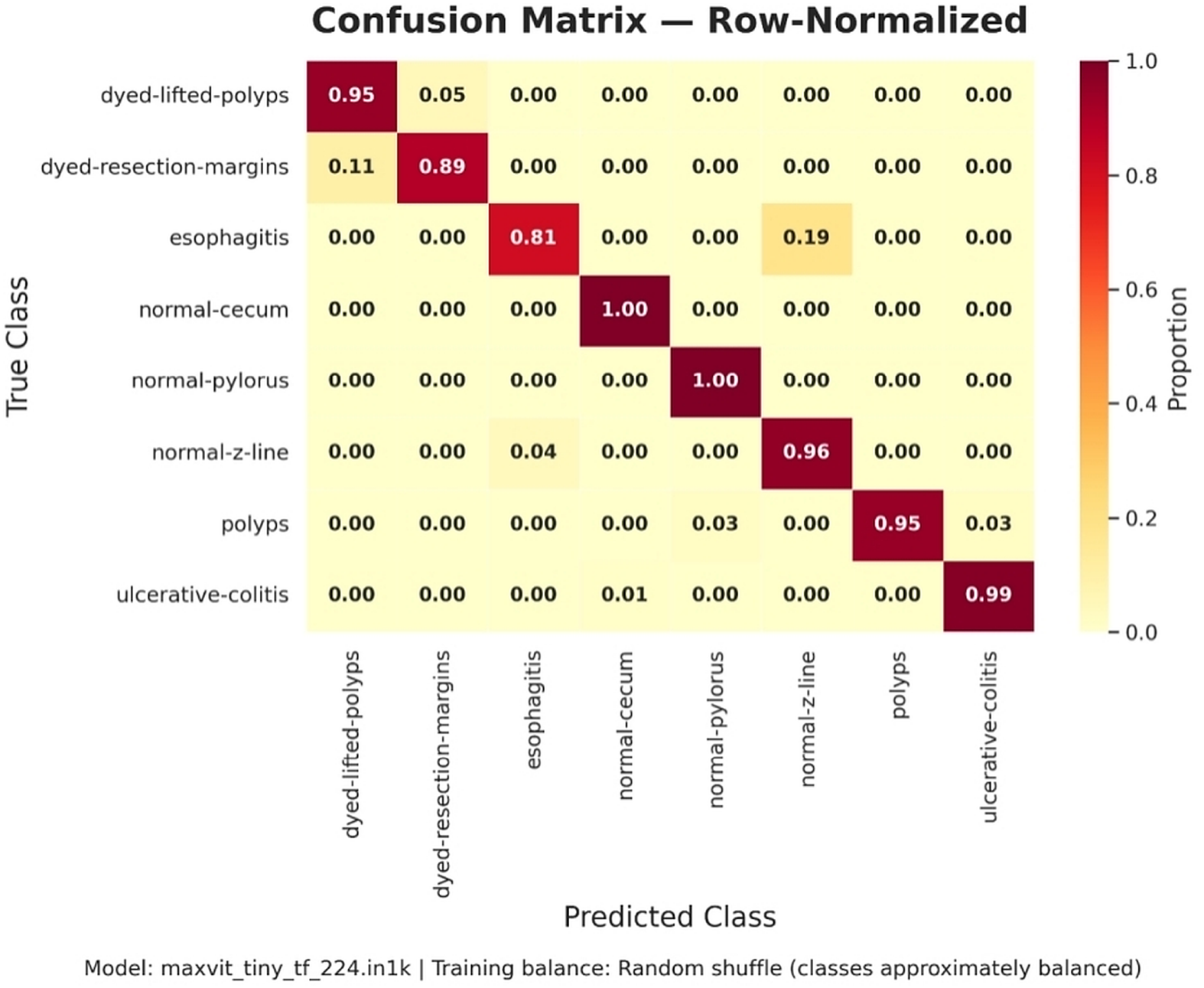
Confusion matrix of the proposed model on the test set (seed 44).

One-vs-rest receiver operating characteristic (ROC) curves for each of the eight classes are shown in Fig 6. The discrimination performance of the model is very high, with per-class AUC values of 0.992–1.000 and a macro-averaged AUC of 0.9975. Near-perfect separability (AUC ≥ 0.9998) was achieved for normal-cecum, normal-pylorus, polyps, and ulcerative-colitis, while dyed-lifted-polyps and dyed-resection-margins yielded AUCs of 0.9974 and 0.9975, respectively. The lowest values occurred for esophagitis (0.993) and normal-z-line (0.992), the two classes that were primarily confused with each other, as shown by the mutual confusion pattern in Fig 5. All curves rose abruptly towards the upper left corner, with the esophagitis curve rising slightly more gradually, and most classes reached a true positive rate above 0.9 at a false positive rate below 0.05. These results suggest that the residual errors are largely due to visually similar classes and to the class-dependent decision threshold, rather than to a lack of discriminative features.

**Fig 6.**
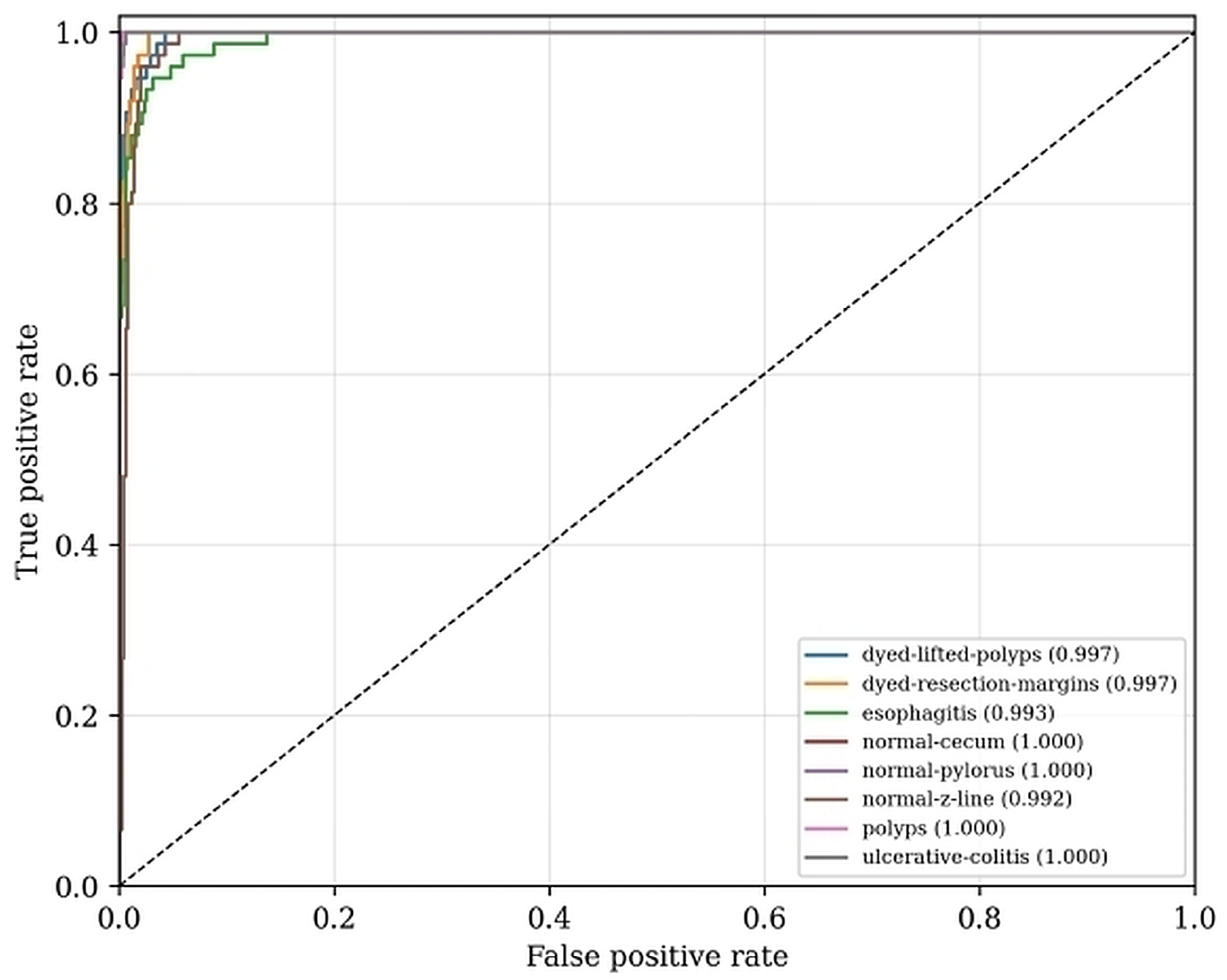
One-vs-rest ROC curves of the proposed model (AUC in the legend).

### Statistical significance analysis

McNemar’s exact test was performed on the paired predictions on the same 600 test images for each of the three seeds (42, 43, and 44) to test whether the difference between the proposed model and each baseline model is statistically significant. Table 6 gives the number of images for which only the proposed model was correct (*b*), for which only the baseline was correct (*c*), and the two-sided *p*-value for each baseline.

**Table 6.**
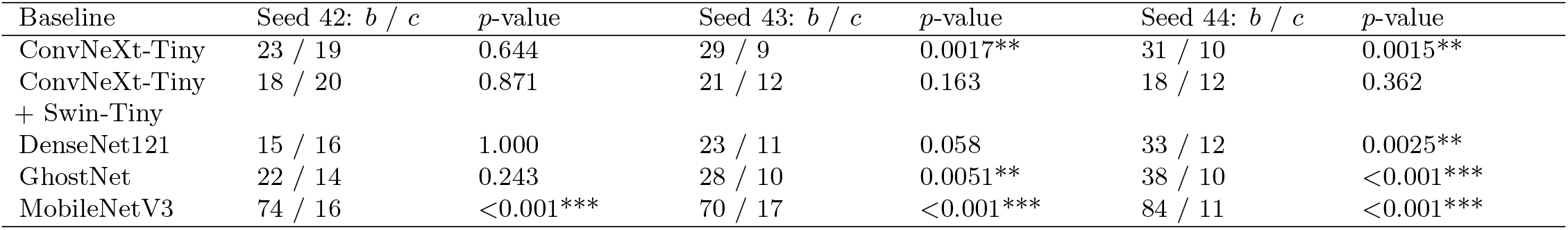
McNemar’s test: proposed MaxViT-Tiny versus baseline models for seeds 42, 43, and 44.

| Baseline | Seed 42: $b / c$ | $p$-value | Seed 43: $b / c$ | $p$-value | Seed 44: $b / c$ | $p$-value |
| --- | --- | --- | --- | --- | --- | --- |
| ConvNeXt-Tiny | 23 / 19 | 0.644 | 29 / 9 | 0.0017** | 31 / 10 | 0.0015** |
| ConvNeXt-Tiny + Swin-Tiny | 18 / 20 | 0.871 | 21 / 12 | 0.163 | 18 / 12 | 0.362 |
| DenseNet121 | 15 / 16 | 1.000 | 23 / 11 | 0.058 | 33 / 12 | 0.0025** |
| GhostNet | 22 / 14 | 0.243 | 28 / 10 | 0.0051** | 38 / 10 | <0.001*** |
| MobileNetV3 | 74 / 16 | <0.001*** | 70 / 17 | <0.001*** | 84 / 11 | <0.001*** |

The proposed model showed a significant improvement over MobileNetV3 in all three seeds (*p <* 0.001). In seeds 43 and 44, it was also significantly better than ConvNeXt-Tiny (*p* = 0.0017 and 0.0015) and GhostNet (*p* = 0.0051 and *p <* 0.001), and in seed 44 it was additionally better than DenseNet121 (*p* = 0.0025). All of these differences are significant at the per-seed Bonferroni criterion of 0.01. At the Bonferroni threshold across all comparisons, 0.05/15 ≈ 0.0033, all differences remain significant except GhostNet in seed 43 (*p* = 0.0051); using the less conservative Holm procedure, all eight significant differences remain. However, the significance was not the same for all seeds: in seed 42 only MobileNetV3 was significantly outperformed by the proposed model, while there was no significant difference against the other models, for example ConvNeXt-Tiny + Swin-Tiny (18 versus 20 discordant images) and DenseNet121 (15 versus 16). No difference against the hybrid was significant in any seed (*p* ≥ 0.163), so this model can be considered statistically comparable to the proposed model on this test set. Two limitations apply. First, because the two trained models are compared on a fixed 600-image test set, McNemar’s test indicates whether the models differ on these images, but not how much of the difference is attributable to training randomness or to a different patient population. Second, all three seeds share the same split and test images, so the three sets of tests are not independent replicates and were not combined into a single *p*-value. The results thus suggest a consistent improvement over MobileNetV3 and a seed-dependent improvement over ConvNeXt-Tiny, GhostNet and DenseNet121.

### Consistency across seeds

The accuracy over the three seeds is summarised in Table 7. The proposed MaxViT-Tiny model achieved the highest mean accuracy (93.72 ± 1.06%) and mean F1-score (0.9370 ± 0.0108) among all models, followed by the ConvNeXt-Tiny + Swin-Tiny hybrid (93.00 ± 0.29%). The lower accuracy in seed 42 (92.50%) compared with seeds 43 and 44 (94.33% in both) accounts for the larger standard deviation of the proposed model.

**Table 7.** Test accuracy (%) per seed and mean *±* standard deviation across seeds.

| Model | Seed 42 | Seed 43 | Seed 44 | Acc. mean ± SD (%) | F1 mean ± SD |
| --- | --- | --- | --- | --- | --- |
| ConvNeXt-Tiny | 91.83 | 91.00 | 90.83 | $91.22 \pm 0.54$ | $0.9120 \pm 0.0054$ |
| ConvNeXt-Tiny + Swin-Tiny | 92.83 | 92.83 | 93.33 | $93.00 \pm 0.29$ | $0.9298 \pm 0.0030$ |
| DenseNet121 | 92.67 | 92.33 | 90.83 | $91.94 \pm 0.98$ | $0.9189 \pm 0.0100$ |
| GhostNet | 91.17 | 91.33 | 89.67 | $90.72 \pm 0.92$ | $0.9064 \pm 0.0092$ |
| MobileNetV3 | 82.83 | 85.50 | 82.17 | $83.50 \pm 1.76$ | $0.8333 \pm 0.0185$ |
| MaxViT-Tiny (Proposed) | 92.50 | 94.33 | 94.33 | $93.72 \pm 1.06$ | $0.9370 \pm 0.0108$ |

The standard deviation is the sample SD ( $n = 3$ seeds).

### Computational cost

The proposed MaxViT-Tiny model has 30.44 M parameters (all trainable), a size of 116.12 MB, and requires 5.33 G MACs (10.66 GFLOPs) for a 224 × 224 image (Table 8). On a Tesla T4 GPU, the inference latency was 32.59 ms per image (30.69 images per second) at batch size 1, which is suitable for offline analysis, but the network is not lightweight compared with mobile architectures such as MobileNetV3 and GhostNet.

**Table 8.** Computational complexity of the proposed model.

| Measure | Value |
| --- | --- |
| Input size | 224 × 224 |
| Parameters (total/trainable) | 30.44 M / 30.44 M |
| Model size | 116.12 MB |
| MACs | 5.33 G |
| FLOPs | 10.66 G |
| Inference latency (batch size 1) | 32.59 ms/image |
| Throughput | 30.69 images/s |
| Device | NVIDIA Tesla T4 |

### Robustness to adversarial attacks

The results of the proposed MaxViT-Tiny model under adversarial attacks are presented in Table 9. The attacks had a major effect on both checkpoints. The clean-trained model achieved a robust accuracy of only 0.33% against the Projected Gradient Descent (PGD) attack, 4.50% against the Carlini–Wagner L2 (CW-L2) attack, and 1.17% against the Momentum Iterative Fast Gradient Sign Method (MI-FGSM). Under the Fast Gradient Sign Method (FGSM), it retained an accuracy of 67.17%.

**Table 9.** Clean and adversarial accuracy on the test set (600 images).

| Checkpoint | Attack | Clean Accuracy (%) | Robust Accuracy (%) | ASR (%) |
| --- | --- | --- | --- | --- |
| Clean-trained | FGSM | 94.33 | 67.17 | 28.80 |
| Clean-trained | PGD | 94.33 | 0.33 | 99.65 |
| Clean-trained | CW-L2 | 94.33 | 4.50 | 95.23 |
| Clean-trained | MI-FGSM (unseen) | 94.33 | 1.17 | 98.76 |
| Adversarially fine-tuned | FGSM | 92.50 | 74.17 | 20.00 |
| Adversarially fine-tuned | PGD | 92.50 | 3.33 | 96.40 |
| Adversarially fine-tuned | CW-L2 | 92.50 | 12.00 | 87.03 |
| Adversarially fine-tuned | MI-FGSM (unseen) | 92.50 | 3.00 | 96.76 |

With adversarial fine-tuning, the accuracy under FGSM improved from 67.17% to 74.17%, and under CW-L2 from 4.50% to 12.00%. The accuracy under PGD and MI-FGSM remained very low, at 3.33% and 3.00%, respectively. These results suggest that single-step adversarial training provides only limited protection against stronger iterative attacks. The adversarially fine-tuned model had a clean accuracy of 92.50%, compared with 94.33% for the clean-trained model, indicating a minor decline in clean-image performance.

Overall, the results show that although the proposed model performs well on clean images, it remains susceptible to adversarial perturbations. Adversarial fine-tuning with FGSM improved robustness against FGSM and CW-L2, but did not defend against PGD or the unseen MI-FGSM attack. Consequently, the proposed model should not be considered adversarially robust without exploring stronger defence approaches, e.g., multi-step adversarial training.

### Attack detection

The accuracy of the attack detector on the clean test images and the images generated by the four attacks is 64.1%, with an AUC of 0.857 (Table 10). After thresholding on the validation data, the false positive rate on clean test images was 4.3%, the precision was 98.1%, and the detection rate on attacked test images was only 56.2%. The detector identified PGD with 83.8% and MI-FGSM with 91.0%, but FGSM and CW-L2 with only 26.5% and 23.8%, respectively. The detector is therefore only a preliminary component of a defence and not a complete defence.

**Table 10.** Performance of the attack detector on the test set.

| Accuracy | Precision | Recall | F1-score | AUC | Balanced acc. |
| --- | --- | --- | --- | --- | --- |
| 0.6410 | 0.9811 | 0.5621 | 0.7147 | 0.8573 | 0.7594 |

### Qualitative explanation

One correctly classified test image for each of the eight classes (prediction confidence 0.854–0.980) is shown with the corresponding explanation maps in Fig 7. In several classes the maps are based on the lesion or the anatomical landmark. For dyed-resection-margins, both Grad-CAM++ and Score-CAM target the resection margin at the edge of the dyed area; for normal-pylorus, both target the pyloric opening; and for polyps, Score-CAM covers the polyp mass. For esophagitis, Score-CAM concentrates on the mucosa next to the lumen, while the Grad-CAM++ map is more scattered, with another peak on the left side of the image. For dyed-lifted-polyps, Grad-CAM++ covers the whole mucosal fold and dyed area, whereas Score-CAM is poor and sparse. In other examples, however, the maps highlight non-anatomical areas: Grad-CAM++ highlights the on-screen annotation box in normal-cecum and polyps, the image border in normal-z-line, and the time-stamp area in ulcerative-colitis, and Score-CAM also weights text overlays in normal-z-line and ulcerative-colitis. This suggests that the network may use these image artefacts to some extent, as they are present in some of the Kvasir images. The figure does not indicate how often this occurs, and these maps are only qualitative evidence, not clinical validation, since only one image per class is displayed. GradientSHAP generated sparse pixel-level attributions that are harder to interpret visually.

**Fig 7.**
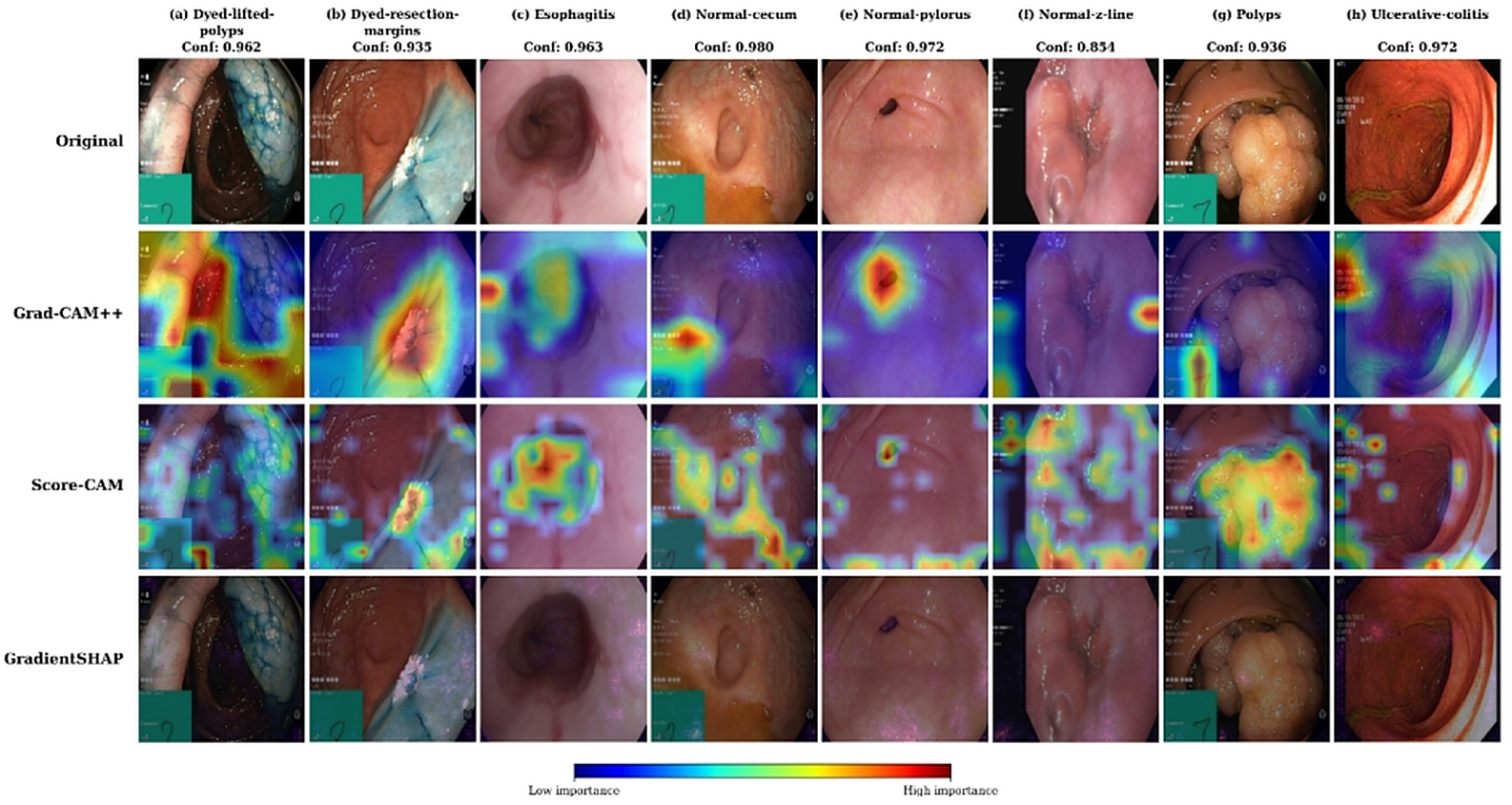
Explanation maps (Grad-CAM++, Score-CAM, GradientSHAP) for one correctly classified test image of each of the eight classes (seed 44; original images from the Kvasir dataset [29]). The class and the prediction confidence are given at the top of each column.

### External validation on GastroVision

Table 11 lists the seven GastroVision classes that were retained, the Kvasir classes they were mapped to, and the number of external images per class.

**Table 11.** Class mapping between GastroVision and Kvasir and the number of external images used per class.

| GastroVision folder | Kvasir class | Images used |
| --- | --- | --- |
| Cecum | normal-cecum | 113 |
| Colon polyps | polyps | 819 |
| Dyed-lifted-polyps | dyed-lifted-polyps | 141 |
| Dyed-resection-margins | dyed-resection-margins | 246 |
| Esophagitis | esophagitis | 107 |
| Gastroesophageal_junction_normal z-line | normal-z-line | 330 |
| Pylorus | normal-pylorus | 393 |
| Total | 7 classes | 2,149 |

The results of the proposed MaxViT-Tiny model on the Kvasir test set (600 images, eight classes) and on the external GastroVision images (2,149 images, seven shared classes) are compared in Table 12. On GastroVision, the model reached an accuracy of 86.23%, a macro F1-score of 0.8025 (95% CI 0.7832 to 0.8233), a Cohen’s kappa of 0.8220, and a macro AUC of 0.9821, compared with 94.33%, 0.9431, 0.9352, and 0.9975 on Kvasir. The accuracy therefore dropped by 8.10 percentage points and the macro F1-score by 14.06 percentage points, while the AUC dropped by only 0.0154. The two evaluations are not exactly like-for-like, since the Kvasir test set has eight balanced classes, while the external set has seven imbalanced classes. The decrease in macro F1-score is larger than that in accuracy because the smaller external classes are not identified to the same extent (Table 13). In the stricter eight-way evaluation the accuracy was 82.18%, and 5.30% of the external images were classified as ulcerative-colitis, a class that is not present in the external data.

**Table 12.** Performance of the proposed model on the Kvasir test set and on the external GastroVision dataset (seed 44).

| Dataset | Acc. (%) | Precision | Recall | F1-score | Kappa | AUC |
| --- | --- | --- | --- | --- | --- | --- |
| Kvasir (test set; 600 images, 8 classes) | 94.33 | 0.9459 | 0.9433 | 0.9431 | 0.9352 | 0.9975 |
| GastroVision (external; 2,149 images, 7 classes) | 86.23 | 0.8213 | 0.7982 | 0.8025 | 0.8220 | 0.9821 |
| Difference (external – Kvasir) | −8.10 | −0.1246 | −0.1451 | −0.1406 | −0.1132 | −0.0154 |

**Table 13.** Class-wise results of the proposed model on the GastroVision external set (seed 44).

| Class | Support | Precision | Recall | F1-score | AUC |
| --- | --- | --- | --- | --- | --- |
| dyed-lifted-polyps | 141 | 0.7834 | 0.8723 | 0.8255 | 0.9852 |
| dyed-resection-margins | 246 | 0.9372 | 0.9106 | 0.9237 | 0.9962 |
| esophagitis | 107 | 0.4662 | 0.5794 | 0.5167 | 0.9622 |
| normal-cecum | 113 | 0.9351 | 0.6372 | 0.7579 | 0.9766 |
| normal-pylorus | 393 | 0.9625 | 0.7837 | 0.8640 | 0.9892 |
| normal-z-line | 330 | 0.7323 | 0.8455 | 0.7848 | 0.9754 |
| polyps | 819 | 0.9323 | 0.9585 | 0.9452 | 0.9900 |
| Macro average | 2,149 | 0.8213 | 0.7982 | 0.8025 | 0.9821 |

The row-normalised confusion matrix of the proposed model is shown in Fig 8. Polyps (0.96), dyed-resection-margins (0.91), and dyed-lifted-polyps (0.87) were recognised best. The same pair of classes that caused most errors on Kvasir (Section 4.4) is also the main problem on the external data: the weakest class was esophagitis (F1-score 0.517, precision 0.466), with 43 of 107 images (40%) predicted as normal-z-line, and 50 of 330 normal-z-line images (15%) predicted as esophagitis. For normal-cecum, the precision was high (0.935), but the recall was low (0.637) because 30% (34/113) of its images were predicted as polyps. Similarly, 14% (56/393) of the normal-pylorus images were predicted as normal-z-line. ROC curves for all classes were high (Fig 9), with AUC between 0.962 and 0.996. This means that the class ranking is well preserved, whereas the decision thresholds learned on Kvasir do not transfer equally well to the new data (recalibration was not tested here). The ECE on GastroVision was 0.039.

**Fig 8.**
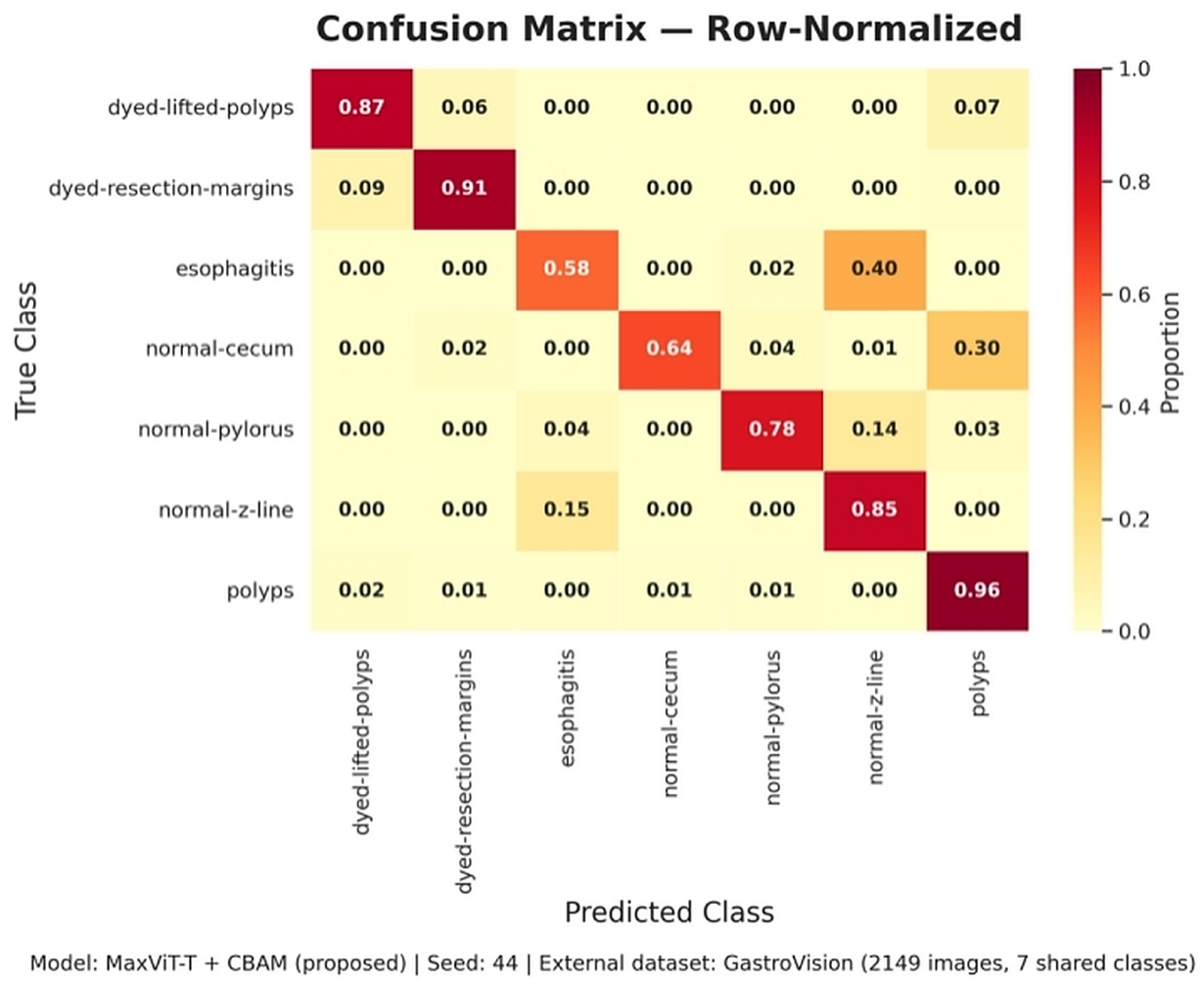
Row-normalised confusion matrix of the proposed model on the GastroVision external set (seed 44, 2,149 images, 7 shared classes).

**Fig 9.**
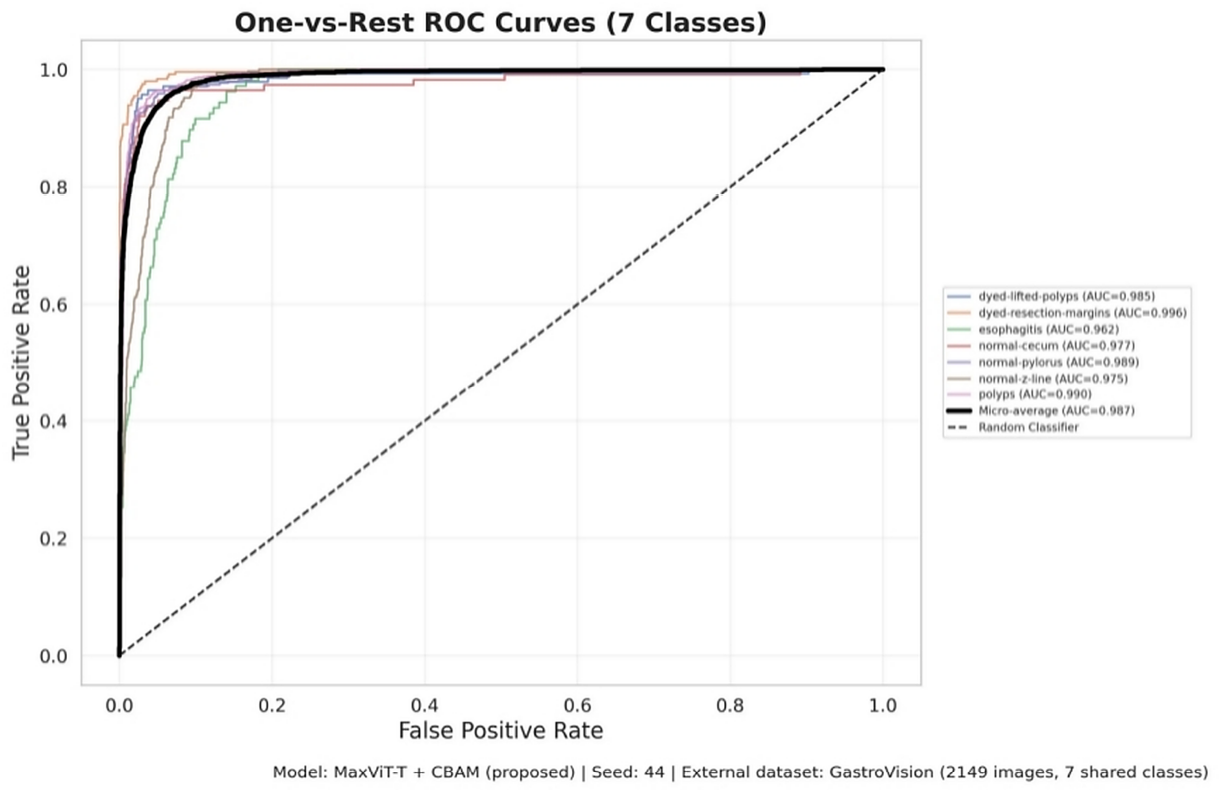
One-vs-rest ROC curves of the proposed model on the GastroVision external set (AUC in the legend).

There are several limitations to this external evaluation. Only one training seed was tested, and external results may differ for other seeds. Only seven of the eight Kvasir classes could be examined, the class mapping was based on folder names, and ulcerative-colitis was assumed to be absent from the external data. The test is not completely source-independent since both Kvasir and GastroVision were partially collected at Bærum Hospital. Finally, the external classes are not well balanced, the smallest classes having 107 and 113 images, so class-wise scores are uncertain. The results show that the performance of the model on an external dataset is still good, although it drops considerably, and the evaluation by itself does not demonstrate robustness of the model across centres or devices.

## Discussion

The proposed MaxViT-Tiny model outperformed all five baselines on all metrics (94.33% accuracy, macro F1 of 0.9431, macro AUC of 0.9975) in the seed-44 comparison, which supports the applicability of a hybrid convolution–transformer backbone for endoscopic classification. However, it showed no significant advantage over its closest baseline, the ConvNeXt-Tiny + Swin-Tiny hybrid, in any seed (only 1.00 percentage point, or 6 images, better in seed 44). It was significantly better than MobileNetV3 in all seeds, but better than ConvNeXt-Tiny, GhostNet, and DenseNet121 in only some seeds. Although duplicate files were removed and every model was tested on the same test set, the accuracy is lower than in previous studies on Kvasir variants and capsule-endoscopy datasets, such as [5] (97.84%) and [9] (98.2%), which often use random image-level partitions that tend to overestimate generalization [14]. The errors occurred mainly between visually similar classes (esophagitis and normal-z-line), and some were due to threshold-based decisions; per-class AUC values were at or above 0.992, indicating that the classes are relatively separable. More importantly, high clean accuracy did not translate into robustness: the accuracy dropped to 0.33% under PGD, single-step adversarial fine-tuning only improved the accuracy against FGSM (and CW-L2), and the preliminary detector identified only 56.2% of attacked images. The model therefore cannot be called robust unless multi-step adversarial training or stronger defence strategies are adopted. In addition, the attention of the network was in some cases on annotation boxes, borders, and time stamps, pointing to partial shortcut learning, which can also hinder cross-dataset generalization [14]. The following limitations should be considered when interpreting these findings: white-box attacks were evaluated at a single perturbation size; the same CBAM head was used for all models, so the effect of the attention mechanism alone cannot be separated; the test set is small (600 images); a single public dataset was used for training and testing; and the split was at image level without patient identifiers. With 30.44 M parameters and 32.59 ms per image, the model can be used for offline analysis, but is not suitable for embedded applications.

## Conclusion

MaxViT-Tiny with a CBAM head was proposed in this study to classify 8-class gastrointestinal endoscopic images on a deduplicated Kvasir dataset with a fixed stratified split and three different seeds. The mean accuracy was 93.72 ± 1.06% across the seeds, statistically significantly higher than that of MobileNetV3 in all seeds, higher than that of ConvNeXt-Tiny, GhostNet and DenseNet121 in some seeds, and statistically comparable to the ConvNeXt-Tiny + Swin-Tiny hybrid. The residual errors were concentrated in visually similar pairs of classes, especially esophagitis and normal-z-line.

The study showed that high clean accuracy is not sufficient. Multi-step attacks were very successful against the model, and single-step adversarial fine-tuning did not substantially reduce their success rate; the preliminary attack detector identified only 56.2% of the attacked images, and the explanation maps revealed that the model used image artefacts. It is therefore important to be aware that this is a research prototype and not a clinically ready tool.

Future directions include (1) patient-level splits, cross-dataset testing and validation at multiple centres or in a prospective setting; (2) multi-step adversarial training and more powerful detection or certified defences; (3) removing image overlays and borders and quantitatively assessing explanations; (4) reducing the model size for real-time and embedded usage; and (5) evaluating the backbones with and without the attention module, repeated over several seeds with paired significance tests, to quantify the contribution of CBAM.

## Acknowledgments

Not applicable.

## Declarations

- **Funding:** No funding was received for this study.
- **Competing interests:** The authors declare no competing financial or non-financial interests.
- **Ethics approval and consent to participate:** Not applicable. This study used only publicly available, de-identified datasets (Kvasir and GastroVision). No new data were collected and no human participants were recruited, so ethical approval and informed consent were not required.
- **Consent for publication:** Not applicable.
- **Data availability:** The datasets analysed in this study are publicly available. The Kvasir dataset (version 1; 4,000 images, eight classes) is available from the Simula Research Laboratory at https://datasets.simula.no/kvasir/. GastroVision (8,000 images, 27 classes) is available from the Open Science Framework at https://osf.io/84e7f/. The copy used in this work was obtained from the Kaggle mirror at https://www.kaggle.com/datasets/orvile/gastrovision-gastrointestinal-disease-detection. No new data were collected for this study.
- **Code availability:** The code used in this study is available from the corresponding author upon reasonable request.
- **Author contribution:** Sadek Al Prince and Israt Fatema Shorna contributed equally to this work. Sadek Al Prince: conceptualization, methodology, software, formal analysis, investigation, and writing—original draft. Israt Fatema Shorna: methodology, validation, investigation, and writing—review and editing. Md Mehedi Hasan, Mohammad Ahanaf Akif, and Aziz Misher Mishu: validation, interpretation, and writing—review and editing; Gazi Mehraj Sakib: writing—review and editing. All authors read and approved the final manuscript.

## Notes

### Competing Interest Statement

The authors have declared no competing interest.

